# Frequent paternal mitochondrial inheritance and rapid haplotype frequency shifts in copepod hybrids

**DOI:** 10.1101/2021.05.10.443310

**Authors:** Jeeyun Lee, Christopher S. Willett

**Affiliations:** University of North Carolina at Chapel Hill, Chapel Hill, NC 27599

**Keywords:** mitochondria, paternal leakage, marine invertebrate, nuclear mitochondrial DNA fragment (NUMT), mitonuclear incompatibility

## Abstract

Mitochondria are assumed to be maternally inherited in most animal species, and this foundational concept has fostered advances in phylogenetics, conservation, and population genetics. Like other animals, mitochondria were thought to be solely maternally inherited in the marine copepod *Tigriopus californicus*, which has served as a useful model for studying mitonuclear interactions, hybrid breakdown, and environmental tolerance. However, we present PCR, Sanger sequencing, and Illumina Nextera sequencing evidence that extensive paternal mitochondrial DNA (mtDNA) transmission is occurring in inter-population hybrids of *T. californicus*. PCR on four types of crosses between three populations (total sample size of 376 F1 individuals) with 20% genome-wide mitochondrial divergence showed 2% to 59% of F1 hybrids with both paternal and maternal mtDNA, where low and high paternal leakage values were found in different cross directions of the same population pairs. Sequencing methods further verified nucleotide similarities between F1 mtDNA and paternal mtDNA sequences. Interestingly, the paternal mtDNA in F1s from some crosses inherited haplotypes that were uncommon in the paternal population. Compared to some previous research on paternal leakage, we employed more rigorous methods to rule out contamination and false detection of paternal mtDNA due to non-functional nuclear mitochondrial DNA fragments. Our results raise the potential that other animal systems thought to only inherit maternal mitochondria may also have paternal leakage, which would then affect the interpretation of past and future population genetics or phylogenetic studies that rely on mitochondria as uniparental markers.

## INTRODUCTION

In most animal taxa, maternal mitochondrial inheritance is widely accepted as the norm, as deviations from this mode of inheritance could have significant evolutionary consequences. If paternal mitochondrial DNA (mtDNA) were also transmitted, there could be lethal genomic conflict due to mitochondrial competition (Hurst and Hamilton 1992; Hastings 1992), zygotes may receive sperm mtDNA that is damaged from deleterious effects associated with heavy respiration (Allen 1996), or there could be an increased propagation of deleterious mitochondrial mutations (Xu 2005). To prevent these potential deleterious effects of paternal mtDNA, animals have evolved various mechanisms to exclude paternal mitochondria, such as ubiquitination or degradation of sperm mitochondria in primates (Sutovsky et al. 1999; Sutovsky et al. 2000; Thompson et al. 2003) and fruit flies (Reilly and Thomas Jr. 1980; DeLuca and O’Farrell 2012).

Despite mechanisms to ensure maternal inheritance of mtDNA, reports of paternal mtDNA in the offspring - termed paternal leakage - have been accumulating in hybrids of various animal species. A subset of examples includes paternal leakage in heterospecific or conspecific hybrids of fruit flies (Kondo et al. 1990; Matsuura et al. 1991; Sherengul et al. 2006; Dokianakis and Ladoukakis 2014), mice (Gyllensten et al. 1991; Shitara et al. 1998), periodical cicadas (Fontaine et al. 2007), ticks (Mastrantonio et al. 2019), and nematodes (Hoolahan et al. 2011; Ross et al. 2016). There are several bivalve species in which mitochondria are doubly inherited from both parents as a norm (Zouros et al. 1994; Gusman et al. 2016). While possible artifacts have not been completely ruled out in all of these latter cases, evidence was consistent with at least some paternal leakage in experimental or natural (Matsuura et al. 1991; Mastrantonio et al. 2019) hybrid populations, along with continued maternal transmission of mtDNA. Some of these studies additionally tested and showed that leaked paternal mtDNA could proliferate to later developmental stages (Sherengul et al. 2006; Fontaine et al. 2007) and persist for multiple generations (Gyllensten et al. 1991). This phenomenon likely has significant consequences for how we understand mitochondria and its applications, because the assumption that mitochondria are only inherited maternally in animals has been the basis for using mtDNA as important molecular markers in phylogenetic inference (Avise et al. 1987), population genetics (Wilson et al. 1985), and conservation (Moritz 1994). Additionally, the presence of paternal mtDNA would impact estimates of genetic diversity and gene flow (Takahata and Maruyama 1981; Chapman et al. 1982; Hoolahan et al. 2011).

The mechanism of frequent paternal leakage and the effects of the phenomena on hybrid fitness are presently unknown. It is speculated that paternal leakage is more commonly found in hybrids compared to pure populations because hybridization may disrupt mechanisms that eliminate paternal mtDNA (White et al. 2008), or because it is easier to experimentally detect paternal mtDNA that is genetically different from the maternal mtDNA (Kaneda et al. 1995; Shitara et al. 1998; Sutovsky et al. 2000). Literature on mitochondrial heteroplasmy, where multiple mitochondrial haplotypes are present in a cell, suggest that certain haplotypes are able to increase in abundance rapidly (Yoneda et al. 1992; Blok et al. 1997; White et al. 1999; Brandstätter et al. 2004), but whether this contributes to increased detection of paternal leakage in hybrids has not been explored. Whether specific paternal mtDNA haplotypes and frequent paternal leakage affect hybrid fitness is also unknown. In several species ranging from seed beetles to centrarchid fishes with highly coadapted mitochondrial and nuclear genomes (Burton et al. 2013), hybrids exhibit decreased fitness due to a breakdown in mito-nuclear coadaptation (Burton and Barreto 2012; Wolff et al. 2014). Paternal leakage may have complicated effects on hybrid fitness in such systems with mitonuclear coadaptation: inherited paternal mtDNA may be beneficial if they interact with the paternal nuclear DNA to restore coadpated functions, or alternatively be harmful if they compete with maternal mtDNA and cause genomic conflict (Hastings 1992; Hurst and Hamilton 1992).

Here we report substantial paternal mtDNA inheritance in inter-population hybrids of the marine copepod *Tigriopus californicus. T. californicus* is a rocky intertidal copepod with many genetically distinct populations along the western coast of North America (Burton 1997; Edmands 2001; Willett and Ladner 2009). This species has previously been thought to solely inherit maternal mitochondria (Burton and Lee 1994), and has been an important model to study the role of mtDNA in inter-population hybrid breakdown (Burton 1990; Edmands 1999; Ellison and Burton 2008; Burton and Barreto 2012; Healy and Burton 2020). MtDNA sequences have diverged in allopatry among populations of *T. californicus*, and several lines of evidence suggest that the mitochondrial and nuclear genomes within each population have coevolved (Rawson and Burton 2002; Barreto and Burton 2013; Barreto et al. 2018). When different populations mate, the hybrids may no longer have a complete set of co-adapted mitochondrial and nuclear products from each parent population. This results in suboptimal cellular functions or lowered fitness, called “mitonuclear incompatibility” (Ellison and Burton 2008; Barreto et al. 2015). Since mitochondrial genes greatly influence the fitness of *T. californicus* hybrids, the presence of paternal mtDNA presents novel evolutionary repercussions to consider, like potential impacts of paternal mtDNA on mitonuclear interaction and hybrid fitness.

In this study, we demonstrate paternal leakage in F1 hybrids of crosses between three genetically and geographically distant *T. californicus* populations in California: Abalone Cove (AB), Santa Cruz (SC), and San Diego (SD). Mitochondrial genome-wide divergence between AB and SC is 20.7%, and between AB and SD is 20.8% (Barreto et al. 2018). These values are much greater than the divergence values for coding regions in the nuclear genome, which are 2.54% between AB and SC, and 2.3% between AB and SD (Pereira et al. 2016). Although *T. californicus* has strikingly high levels of mitochondrial divergence even compared to other species within the subphylum Crustacea (Lefébure et al. 2006), it is still classified as a single species because there are little morphological differences across populations (Monk 1941), no premating isolation between populations (Ganz and Burton 1995; Palmer and Edmands 2000), and inter-population hybrids are viable and fertile (Burton 1990; Edmands 1999). One concerning artifact that could lead to patterns masquerading as paternal leakage are nuclear mitochondrial DNA fragments (NUMTs). NUMTs are non-functional pieces of mtDNA that get incorporated into the nuclear genome, and are typically under 1kb (Gaziev and Shaikhaev 2010). If the paternal nuclear genome in the F1 contained NUMTs, that could result in false detection of paternal mtDNA. One of the most comprehensive ways to exclude NUMTs is to first detect NUMTs via sequence analysis of the genome, avoid those regions for diagnostic PCRs and sequencing, and work with large paternal mtDNA fragments in F1s to avoid the typically short NUMTs. We incorporated multiple molecular techniques to rule out artifacts and method-specific biases for detecting paternal mtDNA in F1 hybrids, quantified the proportion of hybrid individuals with paternal leakage, and examined which paternal mitochondrial haplotypes get passed down.

## MATERIALS AND METHODS

### Sample collection

*Tigriopus californicus* samples were collected in 2013-2014 from Abalone Cove, California (AB: 33°44′16″N, 118°22′31″W), Santa Cruz, CA (SC: 36°56′58″N, 122°02′49″W), and San Diego, CA (SD: 32°45′N, 117°15′W). For all populations, copepods were sampled from multiple pools within an outcrop, where pools are genetically similar and mix over time (Burton and Swisher 1984; Willett and Ladner 2009). Collected populations were kept in 35 parts per thousand (ppt) artificial salt water (Instant Ocean, Spectrum Brands, VA), in an environmental chamber at 20 °C and 12:12 hour light:dark cycle. Copepods were reared in these lab conditions for multiple generations before being used in experiments.

### Inter-population cross set up

*Tigriopus californicus* females can be fertilized only after the final molt into adult copepods (Egloff 1966). Therefore, virgin females from pure populations were obtained either by raising small immature copepodids in individual wells of a 24-well culture plate until maturity or by separating a mating pair with an immature female copepodid - a *T. californicus* male mate guards a female until maturity, and upon mating releases the female (Burton 1985). A virgin female from one population and a male from another population were combined in each well of a new 24-well culture plate for one week, and subsequently the male parent was taken out for DNA extraction, while the female parent was left to produce offspring. DNA was extracted from F1 nauplii, copepodids, and adults, which are different developmental stages in chronological order. Four inter-population crosses were set up, with at least 48 replicates for each: SC female (f) x AB male (m), ABf x SCm, SDf x ABm, and ABf x SDm.

### Individual DNA extraction and standard PCR

DNA was extracted from whole bodies of single copepods by immersing them in 20 uL of lysis buffer consisting of Tris hydrochloride, potassium chloride, proteinase K, and Tween 20 (Hoelzel and Green 1992). The mixture was run in a thermocycler at 65 °C for 1 hour then 99 °C for 15 min. When lysing the copepods, special care was taken to exclude contamination of algae and debris - copepods were put in individual bubbles of freshly made artificial salt water for a few minutes, pipetted onto filter paper, then transferred into lysis buffer using a metal probe sterilized with 95% ethanol.

The resulting DNA was used in polymerase chain reaction (PCR) and agarose gel electrophoresis to detect the presence of inherited paternal mtDNA in hybrid individuals. PCR primers were designed to anneal within the *COB* gene (Figure 1) while avoiding areas with NUMTs. Potential regions of NUMTs were identified by using BLAST of the mitochondrial genome of a population (AB GenBank accession DQ917373; SD accession DQ913891.2) to the entire genome assembly of that population (see supplementary Table S1, Table S2). NUMTs for population SC were not identified because preliminary results showed very few hybrids with inherited paternal SC mtDNA. The forward primers contained two consecutive nucleotides at the 3’ end that were unique to each population (and several other nucleotide mismatches unique to each population along the entire primer), thus were predicted to anneal only to one of the populations in the cross pair. Reverse primers were universal to all three populations (see supplementary Table S3). Numerous control PCR reactions using pure population DNA verified that the forward primer specific to one population did not amplify the mtDNA of a different population. These control reactions also verified that there was no off-target amplification of the nuclear genome. PCR was done with one forward and one reverse primer at a time, and genotyping was done using the Promega Corporation PCR kit (catalog # M8295) or the New England Biolabs PCR kit (cat # M0267) with population-specific thermocycler conditions (see supplementary Table S4). PCR products were run on 1% agarose gel and stained in ethidium bromide, and the population origin of the mtDNA was distinguishable based on band size - 481bp was from AB, 895bp from SC, and 898bp from SD (Figure 1). Only F1s with a distinct band that matched the size of the paternal band were scored as having paternal leakage. To be conservative, F1s with a faint and indistinct band that matched the paternal band size were excluded from the paternal leakage count. While this approach may lead to an underestimate of paternal leakage, it ensures that false positive detection of paternal mtDNA is not included the data.

**Figure 1.**
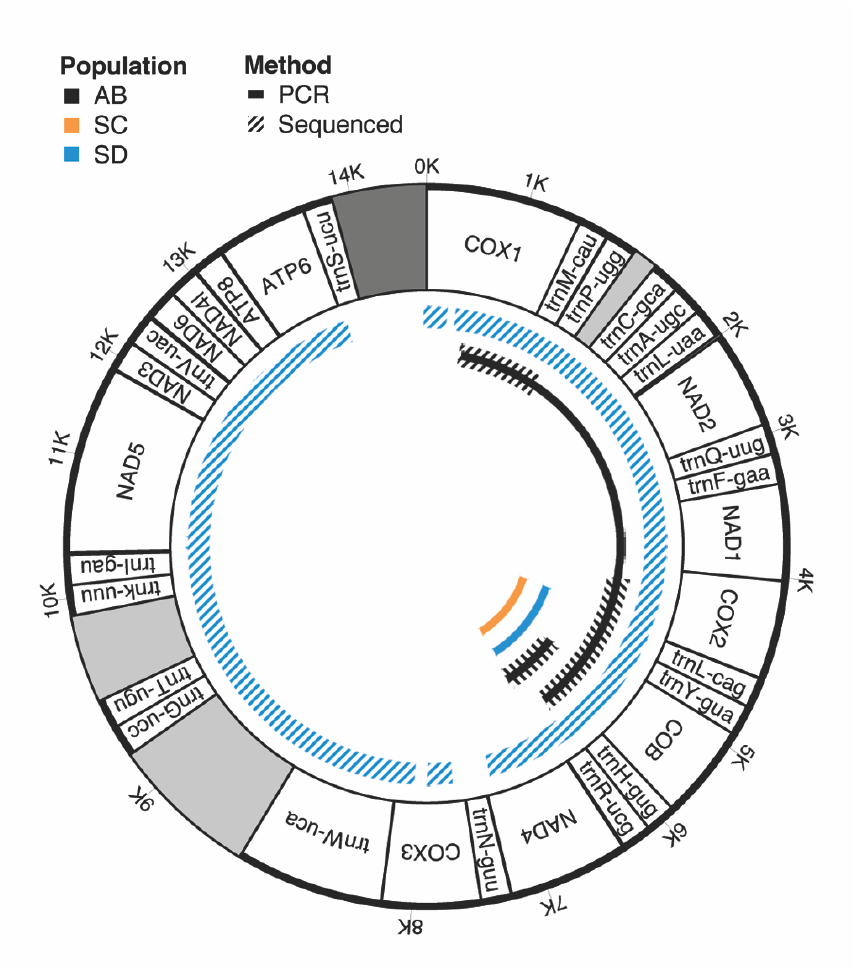
The locations of fragments from PCR (standard PCR and long range PCR) and sequencing methods (Sanger sequencing and Nextera sequencing) along the *Tigriopus californicus* mtDNA (Barreto et al. 2018, accession DQ917373). Different experiments were done for mtDNA from the different *T. californicus* populations AB, SC, and SD. For example, Nextera sequencing was done only for SD along most of the mitochondrial genome, and PCR and Sanger sequencing were done for the short AB fragment in the *COB* gene. Within the gene map, the dark grey region is the control region and the light grey regions are not annotated on NCBI GenBank.

### Plasmid purification of copepod pools and long range PCR

To additionally rule out the possibility of NUMTs, which are generally short and under 1 kb (Gaziev and Shaikhaev 2010), mtDNA was enriched and long range PCR was done to detect long fragments of paternal mtDNA in hybrids of one cross, SCf x ABm. Two pools of approximately 30 AB copepods and 30 SC copepods, each consisting of males, females with and without egg sacs were collected. Another pool of 40 F1s from the cross SCf x ABm, consisting of copepodids, males, and females was collected. MtDNA was enriched for each pool by using Qiagen’s QIAprep Spin Miniprep Kit (catalog # 27104), which aids in purifying plasmid DNA (Quispe-Tintaya et al. 2013).

The mtDNA-enriched DNA was subsequently used in long range PCR. Long range PCR primers were designed to not anneal to NUMT regions, anneal specifically to the AB population and not the SC population (see supplementary Table S3), and span the mitochondrial *COB* gene to *COX1* gene (Figure 1). PCR mixture was prepared using New England Biolabs’ LongAmp® Taq DNA Polymerase (cat # M0323), and run in a thermocycler at 94 °C for 30 sec, cycled 27 times at 94 °C for 30 sec, 59 °C for 15 sec, and 65 °C for 6 min 30 sec, then final extension at 65 °C for 10 min for SC and F1 DNA. For AB paternal population DNA, only 20 cycles were used because the band was too bright to identify the correct fragment size using 27 cycles. Long range PCR products were run on 1% agarose gel then stained in ethidium bromide, and if the sample contained paternal AB mtDNA, large 5-6kb fragments were visible.

### Sanger sequencing of standard PCR, long range PCR products & analysis

PCR products were sequenced using the paternal mtDNA primer to verify similarity to the paternal mtDNA sequence and to inspect if certain haplotypes are more likely to be transmitted to F1s. Specifically, standard PCR products from F1s of SCf x ABm, F1s of SDf x ABm, and the paternal population AB, and long range PCR products from F1s of SCf x ABm and the paternal population AB were purified and Sanger sequenced at Eton Bioscience Inc. using AB-specific primers. The maternal population samples from SD or SC did not produce any PCR bands using AB-specific primers, thus could not be sequenced. For sequencing of long range PCR products, an additional set of primers flanking the *COB* gene (see supplementary Table S3) was used to obtain 2-3kb of sequence.

The sequences were quality trimmed and mapped to the paternal population’s common mitochondrial haplotype in Sequencher (v.5.2.4). The common AB haplotype was obtained by aligning published AB Illumina reads (NCBI SRA SRX2746703) to the published *T. californicus* AB mitochondrial sequence (Burton et al. 2007, GenBank accession DQ917373; Barreto et al. 2018) and extracting the consensus sequence in CLC Genomics Workbench (v.8.0.3). Of note, there were a few possible nucleotide errors in the published mtDNA sequence as there were a number of SNP variants between the published mtDNA and the consensus sequence of the AB Illumina data (see supplementary Table S5). The control region was excluded from the consensus sequence because the coverage was unusually high with lots of repeats, impairing the ability to detect population specific mitochondrial reads. Nucleotide mismatches compared to the most common AB haplotype were identified in sequences of the male parent of the cross and the heteroplasmic F1s. All nucleotide variants were present in the raw pooled AB population Illumina reads.

Further analysis was done to test if all paternal mtDNA haplotypes in the pure AB and F1s of SCf x ABm and SDf x ABm contained any predicted stop codons in their coding regions. First, NCBI blastx was done on the standard PCR and long range PCR sequences of AB to identify the reading frame and coding region. Then the translations of the leaked paternal AB haplotypes in the F1s were compared to that of translations from pure AB individuals in ExPASy (Artimo et al. 2012), to verify the presence/absence of stop codons within the coding region.

### Illumina Nextera sequencing & analysis

A pool of 40 F1s (35 females, 5 males) from cross ABf x SDm was collected for plasmid DNA purification using Qiagen’s QIAprep Spin Miniprep Kit (catalog # 27104). Library was prepped using Illumina’s Nextera™ DNA Flex Library Prep kit and sequencing was done on the NovaSeq 6000.

The sequences were mapped to the common paternal haplotype in CLC Genomics Workbench (v.8.0.3). The common haplotype was obtained by aligning SD Illumina reads to the published *T. californicus* SD mitochondrial sequence and extracting the consensus sequence (Burton et al. 2007; Barreto et al. 2018). Again, the control region was excluded from the consensus sequence. The parameters used for aligning the Nextera reads to the most common AB and SD mtDNA haplotypes were stringent to ensure reads mapped to the correct population’s mtDNA. The parameter settings were as follows: at least 98% of the total alignment length matched the reference sequence, and of that length at least 98% of the identity matched between the aligned read and the reference sequence. Additionally, the cost of mismatch between the read and reference sequence was set to 4 and non-specific matches were ignored; all other parameters were set to default values. This stringent approach may underestimate the true amount of paternal leakage, but provides stronger support for the presence of paternal mtDNA as it minimizes false positive detection of paternal leakage.

### Contamination tests

Several additional procedures were done to verify that paternal mtDNA PCR bands were not a result of laboratory contamination during the DNA lysis and PCR stage. Parts of steps in the individual DNA extraction and PCR procedure mentioned above were purposefully omitted to see if human error could have allowed contamination of paternal mtDNA and resulted in false detection of paternal leakage in the hybrid. A densely-populated AB petri dish was used to obtain DNA or to test for various potential sources of contamination since this was the paternal population for most crosses. In lysis buffer, we put clear water pipetted directly from the dish, water and algae pipetted from the dish (no living copepods were observed under microscope), and soaked a needle tool that was not wiped with ethanol after contacting an AB individual. Replicates of these samples were lysed, PCR amplified using AB-specific primers, and run on a gel along with positive control AB DNA.

## RESULTS

### Frequent paternal leakage in individual F1s and long paternal mtDNA fragment in pooled F1s

Surprisingly, standard PCR of individual F1 hybrids showed frequent paternal leakage for crosses SCf x ABm and SDf x ABm, with 31.1% and 58.5% leakage, respectively (Figure 2). Although not quantitative, the relative brightness of paternal bands in the gel suggested there were at least moderate levels of paternal mtDNA in hybrids of these two inter-population crosses (Figure 3). For the reciprocal crosses ABf x SCm and ABf x SDm, there were fewer F1s with paternal leakage, with 2.0% and 11.7% respectively (Figure 2). The paternal bands in the F1s from both reciprocal crosses were relatively faint compared to the band in the paternal parent, suggesting lower levels of paternal mtDNA (Figure 3). Overall, the reciprocal crosses had fewer individuals showing paternal leakage, and each individual qualitatively appeared to have lower levels of paternal mtDNA. The continued maternal inheritance of mtDNA was verified by the robust PCR amplification of maternal bands in a subset of F1s from all crosses. Doing PCR with primers specific to the paternal population did not show any bands using maternal population samples, indicating that there were no haplotypes in the maternal population that resembled the mitochondrial sequences of the paternal population, and no potential mis-primed sites elsewhere in the maternal genome. Additionally, the control PCRs used to test for laboratory contamination showed no amplified bands on an agarose gel other than the positive control, demonstrating that contamination from the paternal population was unlikely in our samples (see supplementary Figure S1).

**Figure 2.**
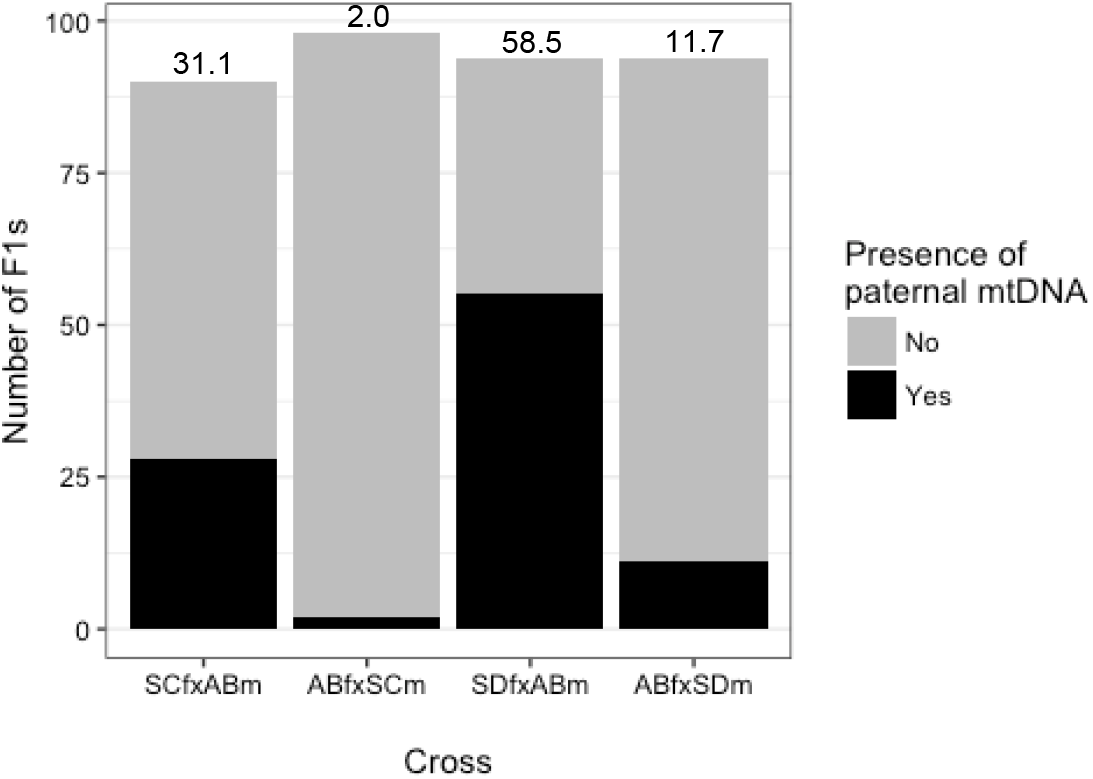
The number of *T. californicus* inter-population F1 offspring with and without paternal leakage based on PCR of the *COB* gene. The percentages of F1s with paternal leakage are indicated at the top of the bars.

**Figure 3.**
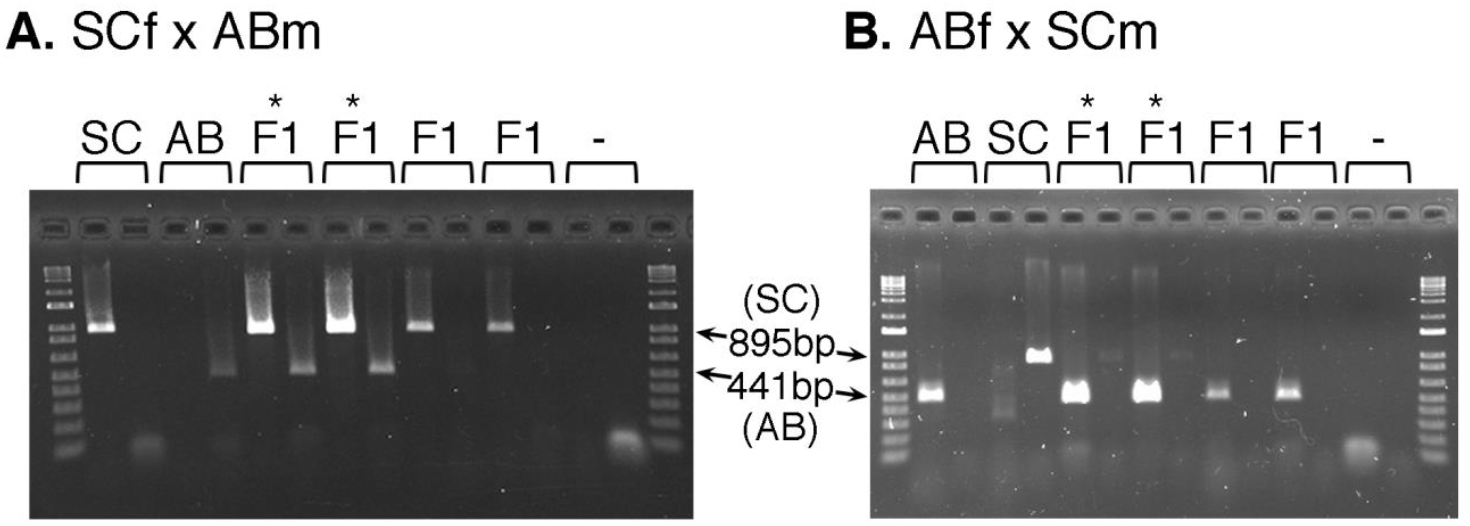
An example of an agarose gel image from *COB* PCR using parental and F1 DNA. The first and last lanes contain the entire range of the Invitrogen™ 1kb Plus DNA ladder (cat # 10787018). The paired lanes show that the same individual’s DNA was used twice for PCR: the first lane of the pair using maternal primers only and the second lane using paternal primers only. DNA from one maternal, one paternal, four F1 individuals, and negative controls consisting of the mastermix from each PCR condition without DNA are shown. F1s with asterisk(*) above have paternal leakage as they show both maternal and paternal mtDNA. **A.** SCf x ABm parents and F1s, and **B.** ABf x SCm parents and F1s.

Long range PCR on F1s of one representative cross type verified that long pieces of paternal mtDNA are present, ruling out short NUMTs (Gaziev and Shaikhaev 2010). Indeed, BLAST results of *T. californicus* showed that the average NUMT length was 223bp with a maximum of 2.4kb in population AB, and an average of 111bp and maximum of 403bp in SD (see supplementary Table S1, Table S2). Long range PCR of the pool of SCf x ABm F1s resulted in a large 5-6kb fragment of AB paternal mtDNA in the agarose gel (Figure 1; see supplementary Figure S2), verifying the presence of paternal mtDNA in F1s as opposed to NUMTs. On the gel, no bands were visible for DNA isolated from the SC maternal population or the PCR negative controls as expected since primers that only amplify AB mtDNA were used.

### Sequencing of leaked paternal fragments reveals haplotype shifts in some crosses

Sanger sequencing of standard and long range PCR products verified that detected fragments in the hybrids are consistent with the paternal mtDNA sequence. Amplified mtDNA sequences in F1s from SCf x ABm and SDf x ABm were highly similar to the most common AB mitochondrial haplotype. For example, in the sequences of the standard PCR products, there were only four nucleotide mismatches out of about 400 bp (*COB* region) in the F1s compared to AB (Table 1). This sequence divergence of 1% shows that there were indeed AB mtDNA in hybrids, particularly when considering the sequence divergence between each paternal population’s *COB* region is around 20%.

**Table 1.** All of the paternal AB haplotype SNP variants in the COB coding region (total length 450-471 bp) in male parents versus F1s with paternal leakage from crosses SCf x ABm and SDf x ABm.

| Cross | Position<sup>a</sup> | 5622 | 5636 | 5747 | 5801 |
| --- | --- | --- | --- | --- | --- |
|  | Nucleotide variants in published AB genome<sup>b</sup> | (T: 98.45%<br>C: 1.55%) | (C: 99.96%<br>A: 0.036%) | (T: 98.18%<br>C: 1.78%<br>A: 0.04%) | (T: 99.72%<br>A: 0.14%<br>C: 0.07%<br>G: 0.07%) |
|  | Common AB haplotype<sup>c</sup> | T | C | T | T |
| SCf x ABm | ABm #1 | T | C | T | T |
|  | ABm #2 | T | T<sup>d</sup> | T | T |
|  | F1 #1 (father ABm #1) | C | C | C | T |
|  | F1 #2 (father ABm #1) | C | C | C | T |
|  | F1 #3 (father ABm #2) | C | C | C | A |
| SDf x ABm | ABm #3 | T | T | T | T |
|  | ABm #4 | T | T | T | T |
|  | F1 #4 (father ABm #3) | C | C | C | T |
|  | F1 #5 (father ABm #3) | C | C | NA<sup>e</sup> | NA |
|  | F1 #6 (father ABm #4) | C | C | C | A |

<sup>a</sup> The nucleotide positions are relative to the reference sequence, Common AB haplotype.
<sup>b</sup> The percentages of nucleotides at each position in the published AB genome containing Illumina reads of a pool of many AB individuals (NCBI SRA SRX2746703).
<sup>c</sup> The reference sequence was the most common AB haplotype, which was obtained by taking the consensus sequence from Illumina reads of many AB individuals (NCBI SRA SRX2746703).
<sup>d</sup> The T variant at position 5636 was not present in the Illumina reads used to obtain the reference sequence, but was present in other published *T. californicus* hybrid sequences (Lima et al. 2019).
<sup>e</sup> Missing nucleotides are indicated with 'NA'.

The four nucleotide mismatches within the *COB* region showed evidence of fast haplotype frequency shifts in the F1 generation (Table 1). At nucleotide positions 5622, 5636, and 5747, the nucleotides in all the F1s were different compared to their fathers in crosses involving both SC and SD females to AB males. At position 5801, a subset of the F1s had different nucleotides compared to the father. Except for the variant at position 5636, all variants in the F1s were present in the Illumina sequence from pooled AB individuals, showing that those sites are heterozygous in the natural paternal population but F1s inherit the uncommon variant. There were no premature stop codons in all paternal haplotypes from AB and F1 individuals (Table 1).

Similarly, Sanger sequencing of long range PCR products showed that the sequence of the amplified band in the SCf x ABm F1s mostly matched the paternal AB mtDNA sequence (Figure 4), but included SNP variants that were uncommon in the AB population. Compared to the most common AB mtDNA haplotype, there were only 11 nucleotide mismatches in the putative paternal mtDNA sequence in pooled F1s, and one mismatch in the pooled ABs (Figure 4; Table 2) out of about 2,250bp. This further corroborates the presence of paternal AB mtDNA in the F1s, and that some haplotypes are common in the F1 but at low frequency in the father individuals or in the general paternal population. Again, all variants in the F1s were found in the general AB population, except for position 5636, where T/C were present in the Sanger sequences while only A/C were present in the Illumina sequences of AB DNA (Table 1; Table 2). To verify that these variants are present in larger pools of AB individuals and not sequencing error, we looked at published hybrid sequences of ABf x SDm F2s (Lima et al. 2019) and found each of the T/A/C variants present at position 5636. Of the 11 SNPs present in the sequenced region of the leaked paternal haplotypes in F1s, only one SNP was predicted to encode a premature stop codon (Table 2).

**Figure 4.**
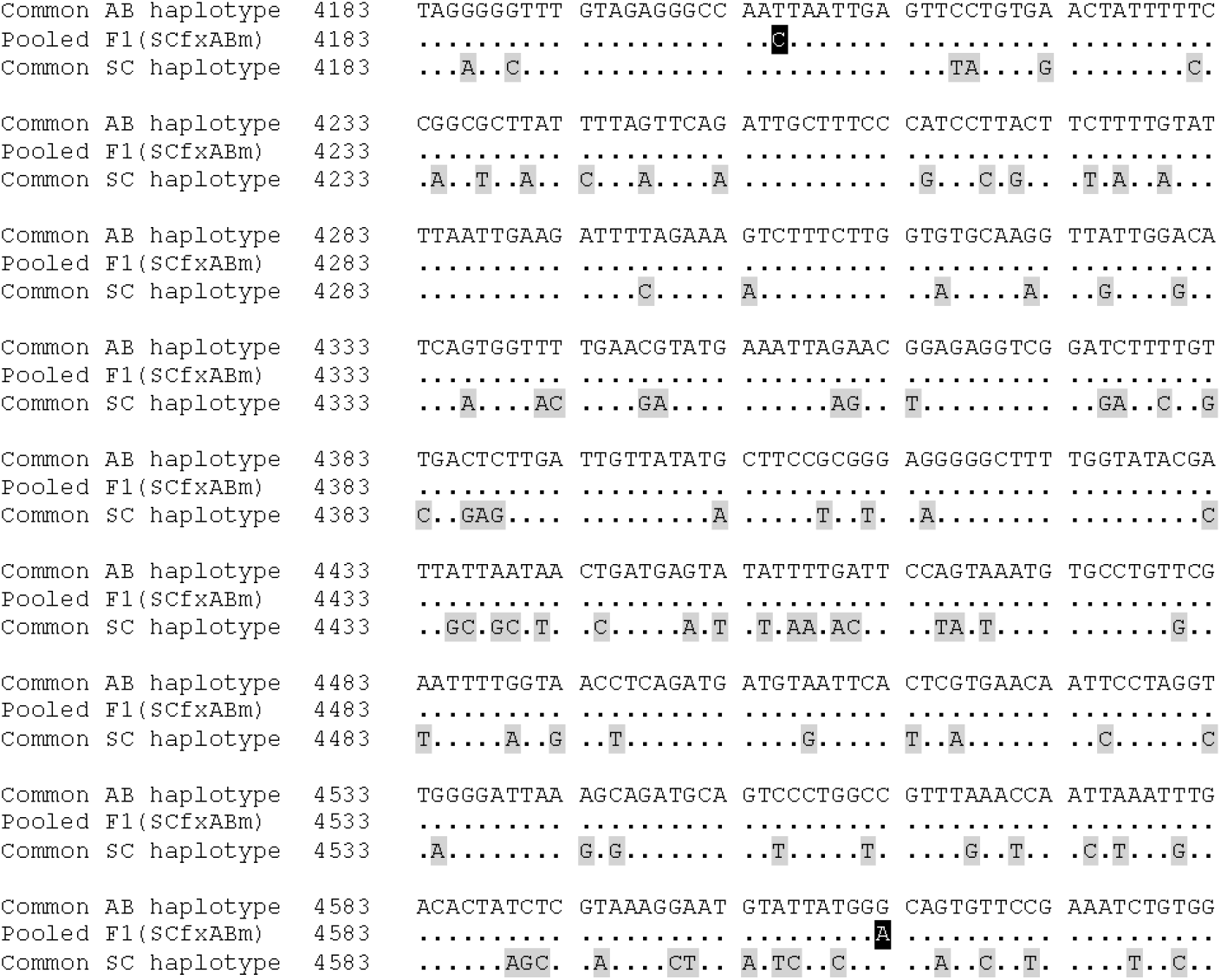
Sanger sequenced long range PCR fragments from plasmid-enriched and pooled SCf x ABm F1s, and the most common SC haplotype aligned to the most common AB haplotype. Only information from nucleotide positions 4183 to 4633 are included as a subsample, and the pooled AB sequence is omitted because it was identical to the reference sequence along this range. SNPs are highlighted with black for the F1s, and grey for the SC sequences, and nucleotides that match the reference are shown with “.”

**Table 2.** All of the paternal AB haplotype nucleotide variants in the Sanger sequence of pooled ABs and pooled SCf x ABm F1s that were used for long range PCR.

|  | AB: 410-1352<sup>a</sup><br>F1: 453-1049 |  |  |  | AB: 4026-4892<br>F1: 4057-4994 |  |  |  |  | AB: 4890-5785<br>F1: 4889-5504 |  |  |  |
| --- | --- | --- | --- | --- | --- | --- | --- | --- | --- | --- | --- | --- | --- |
| Position<sup>b</sup> | 825 | 865 | 971 | 1039 | 4205 | 4612 | 4784 | 4844 | 4849 | 5040 | 5084 | 5622 | 5636 |
| Nucleotide variants (%)<sup>c</sup> | T: 98.86<br>A: 1.11 | T: 99.91<br>G: 0.093 | A: 100 | G: 99.97<br>-<sup>d</sup>: 0.034 | T: 61.12<br>C: 38.85 | G: 99.40<br>A: 0.60 | A: 99.61<br>G: 0.36<br>C: 0.03 | C: 98.53<br>T: 1.47 | T: 98.33<br>C: 1.52 | C: 99.80<br>T: 0.20 | C: 98.64<br>T: 1.31 | T: 98.45<br>C: 1.55 | C: 99.96<br>A: 0.036 |
| Common AB haplotype<sup>e</sup> | T | T | A | G | T | G | A | C | T | C | C | T | C |
| Pooled AB | T | T | A | G | T | G | A | C | T | C | C | T | T/C |
| Pooled F1 (SCf x ABm) | A | G | N<sup>f</sup> | C | C | A/G | G/A | T | C | T<sup>g</sup>/C | T | NA<sup>h</sup> | NA |

<sup>a</sup> There were three sequenced fragments with nucleotide positions indicated in the first row, totaling 2140-2700 bp of sequence.
<sup>b</sup> The nucleotide positions are relative to the reference sequence, Common AB haplotype.
<sup>c</sup> The percentages of nucleotides at each position in the published AB genome containing Illumina reads of a pool of many AB individuals (NCBI SRA SRX2746703).
<sup>d</sup> The symbol '-' indicates there is a deletion in some AB Illumina reads aligned to the common AB haplotype.
<sup>e</sup> The reference sequence was the most common AB haplotype, which was obtained by taking the consensus sequence from Illumina reads of many AB individuals (NCBI SRA SRX2746703).
<sup>f</sup> 'N' corresponds to the IUPAC code for aNy nucleotide, in this case, unknown nucleotide.
<sup>g</sup> The grey shaded cell indicate that T at position 5040 results in a premature stop codon.
<sup>h</sup> 'NA' indicates missing bases because the F1 sequence ended before position 5622.

Nextera sequencing of ABf x SDm F1s showed some fragments that aligned to most of the length of the paternal mitochondrial sequence. Specifically, the Nextera sequences from ABf x SDm F1s covered 84% of the length of the consensus paternal SD mtDNA (Figure 1), albeit with low average coverage of 3.04 and a total of 296 reads. In contrast, when these F1 sequences were mapped to the consensus maternal AB mtDNA, 100% of the reference was covered, with average coverage of 33,448 reads and a total of 3,095,082 reads. Compared to the common haplotype in the general SD population, there were no SNP variants in the F1s’ paternal mtDNA sequences. Standard PCR was done on the same DNA from ABf x SDm F1s used in Nextera sequencing but there was no amplification of SD paternal bands, consistent with very low levels of paternal mtDNA inheritance in this sample (and relatively low in this cross, see Figure 2). Although there is a small amount of paternal mtDNA in these F1s, the presence of fragments that match the majority of paternal mtDNA further decreases the possibility of NUMTs and indicates that the detection of paternal mtDNA is not method-specific.

## DISCUSSION

We found a high prevalence of paternal leakage in the F1 hybrids of *Tigriopus californicus* inter-population crosses, with up to 59% of hybrid individuals from certain crosses having some level of paternal mitochondrial inheritance. We verified the identity of the paternal mtDNA in the hybrids via contamination tests, two methods of PCR and several methods of sequencing. For some crosses, paternal haplotypes that were rare in the paternal population’s mtDNAs were common in the F1s’ mtDNA haplotypes.

The large extent of paternal leakage was unexpected in *T. californicus* hybrids, which were previously thought to only have maternal mitochondrial inheritance (Burton and Lee 1994). While most literature in other species report low copy numbers of paternal mtDNA or less than 10% of hybrids with paternal leakage, there are three studies that show comparable levels as our results for the SCf x ABm and SDf x ABm crosses. In periodical cicada hybrids, there was paternal leakage in 46% of offspring for a particular interspecific cross (Fontaine et al. 2007). There was 19%-48% of paternal leakage in offspring of intraspecific *Drosophila* backcrosses, where the backcross consisted of F1 offspring crossed to a male from the paternal lineage (Sherengul et al. 2006). Additionally, 31%-63% of offspring in interspecific *Drosophila* backcrosses showed paternal leakage (Sherengul et al. 2006). In 40% of hybrids from interpopulation crosses of potato cyst nematodes, the paternal mtDNA replaced the maternal mtDNA (Hoolahan et al. 2011). Reports of frequent paternal leakage may increase if more directions of hybrid crosses are investigated. Several studies showed that reciprocal crosses of the same pair of populations could have contrasting levels of paternal leakage, particularly if one direction of the cross produces fewer or lower fitness offspring (Fontaine et al. 2007; Coleman-Hulbert 2010; Mastrantonio et al. 2019). In the present study, all crosses tested produced roughly 100 offspring (upon which the experiments were stopped) but the reciprocal crosses still showed drastically different prevalence of paternal leakage.

As paternal leakage is unexpected in animals, it is often met with fair criticism that artifacts could be inflating the frequency of paternal mtDNA (Luo et al. 2018; Lutz-Bonengel and Parson 2019; Balciuniene and Balciunas 2019). The previous studies with frequent paternal leakage mentioned above made some efforts to reduce contamination or NUMTs (although Hoolahan et al. (2011) were not explicit about how they did this), but we aimed to make a more thorough attempt to rule out artifacts. Sherengul et al. (2006) avoided NUMTs by amplifying PCR products over 1.8kb, since there were fewer than ten known NUMTs totaling 500bp in *Drosophila melanogaster* at the time (although subsequently greater numbers of NUMTs, some with length 1.5kb, have been found in *D. melanogaster* subgroups) (Rogers and Griffiths-Jones 2012). We took a step further and made new comprehensive lists of NUMTs in relevant *T. californicus* populations, as NUMT discovery is regularly made and NUMTs can be species or population specific (Rogers and Griffiths-Jones 2012). Fontaine et al. (2007) tested for contamination, and minimized NUMTs since they found matching sequences of the *COX1* mitochondrial gene in the hybrids and paternal parent, and verified there were no stop codons in the hybrid paternal sequence. In addition to doing similar procedures to those in Fontaine et al. (2007), we avoided NUMT regions at the primer design stage, and sequencing leaked paternal fragments longer than a single gene, since longer NUMTs are less common.

Other criticisms on reports of paternal leakage question the reliability of the PCR method to detect paternal mtDNA. The present study partially relied on the standard PCR approach for diagnosing paternal mtDNA, which has been widely used in previous literature to confirm paternal leakage (Sherengul et al. 2006, Fontaine et al. 2007, Dokianakis and Ladoukakis 2014, Ladoukakis and Zouros 2017). PCR methods do have several potential issues, for instance, the annealing efficiency of primers may vary across populations or even across haplotypes within a population, causing a bias in the frequency of detected paternal leakage. In the present study, the results from crosses SDfxABm and SCfxABm are still comparable to each other because the same AB population primers were used to diagnose paternal AB mtDNA. It may be that primers for populations SD and SC were worse at annealing compared to AB primers, leading to low amount of paternal leakage in crosses ABf x SDm and ABf x SCm. While the frequency of paternal leakage might be underestimated in these two crosses, it still does not negate our findings that paternal leakage does occur in other cross types of *T. californicus*. Spurious binding of primers was avoided by extensively testing all population primers on many pure population samples before using the primers for detection of paternal mtDNA in hybrids. Another pitfall of PCR could be that mtDNA similar to paternal mtDNA are present in pure maternal samples but were not detected in maternal samples due to experimental error while setting up the PCR reactions. However, the absence of paternal mtDNA bands in maternal individuals was verified using many samples, several PCR machines, multiple PCR trials of positioning the samples in varying locations within the PCR machines, as well as taking other experimental precautions. Moreover, paternal mitochondrial sequences were not found in the Illumina sequences of the pool of maternal population individuals, consistent with the absence of paternal PCR bands. The sequencing results also rule out the possibility of heteroplasmic maternal individuals with haplotypes that are similar to the paternal population’s mtDNA.

Another striking result was that a rare mitochondrial haplotype in the *T. californicus* fathers became the most common paternal haplotype in the F1s of some crosses. If paternal mtDNA were inherited through random sampling of various haplotypes, the most common haplotype in the father would be expected to also be predominant in the offspring. Any deviation in haplotype frequency between generations would indicate some other factor is playing a role in inheritance. In *T. californicus*, it appears that higher frequency and amount of paternal leakage could be correlated with the prevalence of the uncommon paternal haplotype. In our results, 31% of F1s of SCf x ABm had paternal leakage represented by bands almost as intense as paternal control samples on the agarose gel, and had the rare paternal haplotype. Meanwhile, 12% of F1s of ABf x SDm had paternal leakage but showed faint bands, low paternal mtDNA coverage in Nextera sequences, and had the most common SD paternal haplotype.

Previous literature on drastic frequency shifts in maternal mitochondrial haplotypes over a short period of time can give insight into our results. For example, a disease inducing point mutation in the human mitochondria was able to proliferate after one generation (Blok et al. 1997; White et al. 1999), and was even capable of becoming the most common haplotype (Blok et al. 1997). In two independent human families, a point mutation in the mother’s mitochondrial control region became fixed in the child’s mitochondrial genome (Brandstätter et al. 2004). It was speculated that one mitochondrion with the point mutation was preferentially amplified in the oocyte, or that subsets of mtDNA segregated and amplified (Blok et al. 1997; White et al. 1999; Brandstätter et al. 2004). Similar to how specific maternal haplotypes can increase in frequency, particular *T. californicus* paternal haplotypes could be preferentially amplified in the hybrid if they confer replication advantage - for instance a certain paternal haplotype will be amplified more if it is shorter than another haplotype due to a deletion (Yoneda et al. 1992). The premature stop codon in one leaked paternal haplotype in our study (Table 2) indicates some disruption of function, but we don’t know if this stop codon results in large deletions that confer a replicative advantage. For future work, comparing the genetic structures of the various paternal haplotypes would give insight on how haplotype frequencies can shift dramatically.

Several studies provide clues about the mechanism of paternal mtDNA transmission in hybrids. In water frogs, a female hybrid with paternal mtDNA could pass those copies to the offspring (Radojičić et al. 2015), showing that paternal mtDNA can be passed through several generations. Paternal leakage was also reported in *Drosophila* and nematode backcross offspring (Sherengul et al. 2006; Ross et al. 2016). These studies suggest that either paternal mtDNA can be transmitted beyond the F1 stage, or that more opportunities to cross with the paternal lineage increase paternal leakage. Other researchers have speculated on how paternal mtDNA can enter the germline. Ladoukakis and Zouros (2017) suggest that an embryo could contain paternal mtDNA if the egg is genetically divergent from the sperm and fails to recognize and destruct sperm mitochondria during fertilization – this would explain the process of paternal leakage in hybrid individuals. Another insight can be gained by considering mussel species in the family Mytilidae, which, unlike other animal species, have one mitochondrial genome that is inherited from the mother for all offspring and one mitochondrial genome that is inherited from the father but is only found in male offspring (Zouros 2000). Recent studies found that the inherited paternal mtDNA have palindromic motifs in the control region that may increase transcription, have different open reading frames compared to maternal mtDNA, and have an extra cytochrome c oxidase subunit II gene (Passamonti et al. 2011; Zouros 2013). These genetic structures may help explain why transmission of paternal mtDNA along with maternal mtDNA is the norm in these mussels. In the *T. californicus* system, we do not know if there are such sequence differences in the leaked paternal mtDNA. In fact, the exact molecular mechanisms that cause paternal leakage in a typical animal system have not yet been directly tested, and the processes of paternal leakage are much less characterized compared to processes of uniparental maternal mtDNA inheritance.

It is largely unknown how extensive paternal leakage could affect hybrid fitness, but studies on how high amounts of heteroplasmy affect fitness can provide a clue. For example, induced heteroplasmic mice with initially equal proportion of two mitochondrial haplotypes had decreased cognition and physiological functions compared to homoplasmic mice (Sharpley et al. 2012). Harmful point mutations that cause MELAS syndrome in humans were selected against if they contributed more than 80% of the mitochondrial genome relative to the wild type mtDNA (Chomyn et al. 1992; Wallace 1999). In highly heteroplasmic *Drosophila melanogaster* lines, mutant mitochondrial alleles were able to persist if there was another allele that complemented the wild type function (Ma et al. 2014). These studies suggest that high frequency of heteroplasmy can decrease or sometimes have no effect on fitness. It may be that paternal mtDNA in the hybrids behave like mutant mitochondrial haplotypes and thus affect fitness in similar ways. In our study, there is evidence of a deleterious mutation in a leaked paternal haplotype, but it is unclear how this will impact fitness.

Our results challenge a variety of animal studies that utilize mitochondria as molecular markers (Hoolahan et al. 2011), and invite further studies on the prevalence and mechanisms of paternal leakage. Within the *T. californicus* system, future work should involve looking at sequence differences of the uncommon versus common paternal haplotype in the F1 hybrids and quantifying the amount of paternal mtDNA within a hybrid individual. As the process of paternal leakage is demystified, an interesting follow-up evolutionary question would be how extensive paternal leakage affects hybrid fitness. Our findings here will add to the growing evidence of paternal leakage in animals, and spark discussion on how paternal leakage could affect studies on molecular biology and evolution.

## FUNDING

This work was supported by the National Science Foundation’s Division of Integrative Organismal Systems (grant numbers NSF IOS-1155325 to C.S.W., NSF IOS-1555959 to J. Kingsolver and C.S.W.); Kenan Trust Fund; and University of North Carolina at Chapel Hill’s Graduate School Summer Research Fellowship.

## ACKNOWLEDGEMENTS

We thank the “Illumina Workshop In A Box with Nextera Flex @ UNC with HTSF” workshop for Nextera sequencing supplies and services. We also thank Liana Kostak for help with rearing the hybrid crosses and collecting DNA, Jorge Santana, Sarah Slay, MaryAnn Bowyer for help with some of the research, members of Dr. Joel Kingsolver and Dr. Christopher Willett labs who provided feedback on the experimental design and manuscript, and Dr. Maria Servedio and Dr. Daniel Matute for input on the overall experimental design.

## Notes

### Competing Interest Statement

The authors have declared no competing interest.

