## Supplemental Tables and Figures for "Frequent paternal mitochondrial inheritance and rapid haplotype frequency shifts in copepod hybrids"

SUPPLEMENTARY MATERIALS

**Table S1:** BLAST High Scoring Pairs table from CLC Genomics Workbench of NUMTs in population AB (SRX2746703). Sequences were assembled[a] using SPAdes (v. 3.7.0; Bankevich et al. 2012).

| Hit | E-value | HSP start | HSP end | HSP length | Query start | Query end | % Identity | % Gaps |
|---|---|---|---|---|---|---|---|---|
| NODE_17498_length_1815_cov_385.626_ID_3334433 | 0.00 | 1374 | 1 | 1,374.00 | 1 | 1378 | 96.01 | 0.29 |
| NODE_7129_length_6300_cov_39.1425_ID_2181513 | 7.19E-05 | 5776 | 5813 | 38 | 1 | 38 | 92.11 | 0 |
| NODE_9575_length_4271_cov_33.4227_ID_2206176 | 7.19E-05 | 3780 | 3743 | 38 | 1 | 38 | 92.11 | 0 |
| NODE_109363_length_245_cov_2.83158_ID_5069043 | 2.68E-29 | 21 | 190 | 170 | 136 | 305 | 77.65 | 0 |
| NODE_3167_length_15430_cov_30.5033_ID_5435208 | 0.45 | 3480 | 3453 | 28 | 411 | 438 | 92.86 | 0 |
| NODE_31941_length_827_cov_2.12824_ID_3489777 | 6.28E-69 | 23 | 633 | 611 | 537 | 1151 | 69.86 | 4.47 |
| NODE_101907_length_257_cov_1.09901_ID_5001424 | 1.58 | 35 | 14 | 22 | 566 | 587 | 100 | 0 |
| NODE_849_length_39239_cov_28.6111_ID_1473387 | 8.19E-17 | 39239 | 39185 | 55 | 612 | 666 | 98.18 | 0 |
| NODE_16439_length_1985_cov_33.7394_ID_5135620 | 2.68E-10 | 1 | 43 | 43 | 625 | 667 | 97.67 | 0 |
| NODE_6_length_178655_cov_9.29977_ID_1365775 | 8.75E-04 | 61724 | 61761 | 38 | 685 | 722 | 89.47 | 0 |
| NODE_35113_length_741_cov_4.04082_ID_3627594 | 3.26E-09 | 4 | 76 | 73 | 689 | 761 | 80.82 | 0 |
| NODE_11889_length_3164_cov_258.377_ID_4638268 | 8.18E-55 | 56 | 205 | 150 | 1417 | 1566 | 92.67 | 0 |
| NODE_11889_length_3164_cov_258.377_ID_4638268 | 0.00 | 205 | 2582 | 2,378.00 | 1934 | 4317 | 93.71 | 0.34 |
| NODE_69185_length_354_cov_19.0301_ID_6789860 | 4.81E-159 | 1 | 354 | 354 | 2000 | 2353 | 95.48 | 0 |
| NODE_152743_length_181_cov_18.1905_ID_6800962 | 2.34E-74 | 181 | 1 | 181 | 2482 | 2662 | 95.03 | 0 |
| NODE_9265_length_4461_cov_41.493_ID_2155715 | 1.30E-20 | 1449 | 1320 | 130 | 2550 | 2680 | 78.63 | 0.76 |
| NODE_111303_length_243_cov_23.9309_ID_6794938 | 7.64E-106 | 1 | 243 | 243 | 2713 | 2954 | 95.88 | 0.41 |
| NODE_12003_length_3125_cov_2.14039_ID_2351327 | 5.52 | 114 | 151 | 38 | 2955 | 2991 | 84.21 | 2.63 |
| NODE_51654_length_480_cov_24.0353_ID_4399515 | 0 | 1 | 480 | 480 | 3158 | 3636 | 95.83 | 0.21 |
| NODE_73662_length_334_cov_97.7814_ID_4758666 | 2.84E-149 | 334 | 1 | 334 | 4002 | 4335 | 95.51 | 0 |
| NODE_14644_length_2343_cov_2.16783_ID_2543674 | 3.06E-03 | 1 | 30 | 30 | 4125 | 4154 | 96.67 | 0 |
| NODE_155_length_79164_cov_30.6294_ID_4976243 | 9.35E-10 | 3617 | 3561 | 57 | 4272 | 4328 | 87.72 | 0 |
| NODE_216029_length_108_cov_18.6415_ID_5900746 | 1.93E-18 | 55 | 1 | 55 | 4281 | 4335 | 100 | 0 |
| NODE_161117_length_159_cov_1875.56_ID_6154849 | 0.01 | 85 | 57 | 29 | 4396 | 4424 | 96.55 | 0 |
| NODE_9382_length_4393_cov_41.2211_ID_2296750 | 3.97E-08 | 4227 | 4107 | 121 | 4811 | 4931 | 71.9 | 0 |
| NODE_40647_length_628_cov_2.12565_ID_3857435 | 1.93E-18 | 628 | 574 | 55 | 4951 | 5005 | 100 | 0 |
| NODE_100537_length_260_cov_22.9171_ID_6830596 | 6.70E-113 | 260 | 1 | 260 | 5434 | 5694 | 95.4 | 0.38 |
| NODE_4362_length_11127_cov_38.1195_ID_1722301 | 5.52 | 9240 | 9215 | 26 | 5667 | 5692 | 92.31 | 0 |

| | | | | | | | | |
|---|---|---|---|---|---|---|---|---|
| NODE_93133_length_275_cov_26.3227_ID_6808262 | 1.38E-121 | 275 | 1 | 275 | 5821 | 6095 | 95.64 | 0 |
| NODE_65461_length_374_cov_20.4671_ID_6794160 | 1.21E-166 | 374 | 1 | 374 | 6051 | 6425 | 95.2 | 0.27 |
| NODE_117574_length_235_cov_27.0778_ID_6794290 | 3.71E-59 | 1 | 135 | 135 | 6069 | 6203 | 98.52 | 0 |
| NODE_214025_length_110_cov_22.8727_ID_6818316 | 9.98E-16 | 1 | 64 | 64 | 6212 | 6276 | 92.31 | 1.54 |
| NODE_236860_length_84_cov_93.5862_ID_6829876 | 3.72E-21 | 1 | 84 | 84 | 6456 | 6539 | 89.53 | 4.65 |
| NODE_37846_length_680_cov_603.31_ID_4238329 | 0.00 | 680 | 1 | 680 | 6485 | 7163 | 97.94 | 0.15 |
| NODE_38440_length_669_cov_1.3013_ID_3770039 | 4.24E-14 | 669 | 615 | 55 | 6699 | 6753 | 94.55 | 0 |
| NODE_6174_length_7507_cov_108.253_ID_2481257 | 3.24E-180 | 7150 | 7507 | 358 | 6806 | 7163 | 99.44 | 0 |
| NODE_59700_length_411_cov_1.19944_ID_4467868 | 5.52E-19 | 411 | 349 | 63 | 7414 | 7474 | 96.83 | 3.17 |
| NODE_8440_length_5057_cov_46.8699_ID_2098344 | 1.58 | 4167 | 4141 | 27 | 7461 | 7487 | 92.59 | 0 |
| NODE_59834_length_410_cov_2.36901_ID_4470810 | 2.68E-10 | 40 | 1 | 40 | 7568 | 7607 | 100 | 0 |
| NODE_2015_length_22541_cov_30.0636_ID_1527945 | 4.24E-14 | 2238 | 2287 | 50 | 7774 | 7823 | 98 | 0 |
| NODE_12310_length_3003_cov_1.93385_ID_2447844 | 2.51E-23 | 2681 | 2608 | 74 | 8399 | 8472 | 94.59 | 0 |
| NODE_9265_length_4461_cov_41.493_ID_2155715 | 2.86E-16 | 1534 | 1441 | 94 | 8428 | 8512 | 80.85 | 9.57 |
| NODE_497_length_50577_cov_33.3443_ID_3673816 | 5.88E-82 | 97 | 618 | 522 | 8557 | 9096 | 73.54 | 6.2 |
| NODE_1330_length_30356_cov_28.0854_ID_2681346 | 0.45 | 27491 | 27520 | 30 | 8812 | 8843 | 93.75 | 6.25 |
| NODE_39983_length_640_cov_7.73675_ID_3830031 | 2.20E-30 | 1 | 123 | 123 | 8995 | 9118 | 85.48 | 0.81 |
| NODE_30_length_117063_cov_28.7922_ID_1376829 | 1.69E-06 | 44200 | 44155 | 46 | 9127 | 9172 | 89.13 | 0 |
| NODE_18184_length_1723_cov_22.6691_ID_2931649 | 5.52 | 872 | 916 | 45 | 9398 | 9446 | 79.59 | 8.16 |
| NODE_16056_length_2050_cov_32.012_ID_3587190 | 1.58E-19 | 1988 | 2050 | 63 | 10079 | 10142 | 96.88 | 1.56 |
| NODE_21061_length_1410_cov_120.552_ID_4162828 | 0.00 | 1 | 1410 | 1,410.00 | 10425 | 11834 | 94.04 | 0 |
| NODE_286_length_63443_cov_27.9106_ID_1396279 | 1.93E-18 | 63389 | 63443 | 55 | 10628 | 10682 | 100 | 0 |
| NODE_510_length_49922_cov_29.0035_ID_1656637 | 1.93E-18 | 1 | 55 | 55 | 11211 | 11265 | 100 | 0 |
| NODE_9265_length_4461_cov_41.493_ID_2155715 | 4.24E-52 | 1317 | 1082 | 236 | 11303 | 11539 | 80.17 | 0.42 |
| NODE_100019_length_261_cov_22.6214_ID_6814154 | 8.71E-118 | 261 | 1 | 261 | 11842 | 12102 | 96.55 | 0 |
| NODE_6913_length_6538_cov_49.5804_ID_6363637 | 0.45 | 5206 | 5233 | 28 | 11989 | 12016 | 92.86 | 0 |
| NODE_219120_length_105_cov_19.98_ID_6805438 | 1.93E-18 | 105 | 48 | 58 | 12204 | 12261 | 98.28 | 0 |
| NODE_5279_length_9046_cov_29.4813_ID_1808098 | 5.52 | 4390 | 4424 | 35 | 12224 | 12256 | 85.71 | 5.71 |
| NODE_34746_length_750_cov_322.86_ID_3905349 | 0.00 | 94 | 750 | 657 | 12257 | 12913 | 95.43 | 0 |
| NODE_214025_length_110_cov_22.8727_ID_6818316 | 1.93E-18 | 56 | 110 | 55 | 12433 | 12487 | 100 | 0 |
| NODE_643_length_45064_cov_39.6533_ID_1465119 | 5.52 | 4139 | 4174 | 36 | 12996 | 13031 | 83.33 | 0 |
| NODE_216095_length_108_cov_20.2264_ID_6820838 | 1.93E-18 | 108 | 54 | 55 | 13142 | 13196 | 100 | 0 |
| NODE_137293_length_214_cov_0.823899_ID_5262525 | 0.45 | 62 | 16 | 47 | 13711 | 13755 | 82.98 | 4.26 |

[a] The published genomes for the AB populations of *T. californicus* (Barreto et al. 2018) were constructed based on mapping sequences from this population to the SD population's relatively high quality genome and therefore do not contain unique sequences not found in the SD genome (potentially including unique mtDNA pseudogenes). Lower quality de novo assemblies were made from the AB population in an attempt to uncover novel insertions of mtDNA into the nuclear genomes of this population. The Illumina 400bp sequences from the AB population (NCI SRA SRX2746703) were run on the computational cluster at UNC using the default parameters of SPAdes (v. 3.7.0; Bankevich et al. 2012). The resulting AB genome was highly fragmented and contained a number of bacterial contaminant scaffolds. To help remove some of these contaminating sequences, reads were mapped back to assemblies using BWA-MEM (Li and Durbin 2009) and the coverage depth calculated for each scaffold using the idxstats command in SAMtools. For the resulting AB assembly, the modal coverage was 70 and scaffolds were trimmed as follows: for scaffolds >10kb those with coverage less than 40 were removed (most were checked by BLAST searches and confirmed to be bacterial); for scaffolds between 10kb and 1kb removed scaffolds with coverages less than 20; for scaffolds between 1kb and 400bp trimmed out those with coverage less than 10; and finally all scaffolds less than 400bp size were removed. This trimming resulted in a final assembly size of 182Mb for AB, which is comparable in size to the initial assembly of the SD genome (NCBI SRA SRX469409) with bacterial contaminants removed of 181Mb and the improved published assembly size of 191Mb.

**Table S2.** BLAST High Scoring Pairs table from CLC Genomics Workbench of NUMTs in the published genome of population SD (genome v.2.1; SRX469409).

| Hit | E-value | HSP start | HSP end | HSP length | Query start | Query end | %Identity | %Gaps |
|---|---|---|---|---|---|---|---|---|
| Chromosome_12 | 2.39E-03 | 15239773 | 15239819 | 47 | 79 | 125 | 82.98 | 0 |
| Chromosome_8 | 4.32 | 5082335 | 5082383 | 49 | 603 | 651 | 77.55 | 0 |
| Chromosome_9 | 6.83E-23 | 15359603 | 15359538 | 66 | 606 | 671 | 98.48 | 0 |
| Chromosome_10 | 3.77E-102 | 6619264 | 6618870 | 395 | 853 | 1248 | 81.31 | 0.25 |
| Chromosome_12 | 4.32 | 5899001 | 5899029 | 29 | 1185 | 1213 | 89.66 | 0 |
| Chromosome_6 | 3.54E-39 | 2814373 | 2814186 | 188 | 1201 | 1388 | 80.42 | 1.06 |
| Chromosome_6 | 4.32 | 14915676 | 14915651 | 26 | 1574 | 1599 | 92.31 | 0 |
| Chromosome_6 | 4.32 | 14486695 | 14486660 | 36 | 1578 | 1613 | 83.33 | 0 |
| Chromosome_10 | 4.32 | 12407921 | 12407893 | 29 | 1646 | 1674 | 89.66 | 0 |
| Chromosome_10 | 4.32 | 5519762 | 5519792 | 31 | 1652 | 1682 | 87.1 | 0 |
| Chromosome_1 | 8.85E-123 | 580133 | 580379 | 247 | 1724 | 1970 | 100 | 0 |
| Chromosome_4 | 8.90E-09 | 8781406 | 8781480 | 75 | 1726 | 1799 | 81.58 | 3.95 |
| Chromosome_7 | 4.32 | 10252930 | 10252907 | 24 | 1909 | 1932 | 95.83 | 0 |
| Chromosome_10 | 4.32 | 10183088 | 10183063 | 26 | 2154 | 2179 | 92.31 | 0 |
| Chromosome_1 | 1.24 | 11022735 | 11022761 | 27 | 2631 | 2657 | 92.59 | 0 |
| Chromosome_8 | 4.32 | 3307481 | 3307513 | 33 | 3006 | 3036 | 87.88 | 6.06 |
| Chromosome_8 | 4.32 | 9952271 | 9952302 | 32 | 3590 | 3624 | 88.57 | 8.57 |
| Chromosome_9 | 2.39E-22 | 8682487 | 8682628 | 142 | 3619 | 3760 | 77.46 | 0 |
| Chromosome_9 | 1.24 | 15543257 | 15543288 | 32 | 3868 | 3899 | 87.5 | 0 |
| Chromosome_11 | 1.24 | 12919553 | 12919520 | 34 | 4128 | 4160 | 88.24 | 2.94 |
| Chromosome_12 | 0.35 | 1251536 | 1251514 | 23 | 4213 | 4235 | 100 | 0 |
| Chromosome_11 | 2.23E-16 | 15770495 | 15770419 | 77 | 4240 | 4313 | 87.01 | 3.9 |
| Chromosome_1 | 4.32 | 1582043 | 1582083 | 41 | 4595 | 4634 | 82.93 | 2.44 |
| Chromosome_12 | 2.90E-40 | 3286787 | 3287103 | 317 | 4624 | 4941 | 73.46 | 4.01 |
| Chromosome_8 | 4.32 | 2706877 | 2706900 | 24 | 4928 | 4951 | 95.83 | 0 |
| Chromosome_6 | 4.32 | 345575 | 345600 | 26 | 6285 | 6310 | 92.31 | 0 |
| Chromosome_8 | 4.32 | 148306 | 148281 | 26 | 7326 | 7351 | 92.31 | 0 |
| Chromosome_7 | 4.32 | 7228337 | 7228380 | 44 | 7812 | 7855 | 79.55 | 0 |
| Chromosome_10 | 0.35 | 1910944 | 1910981 | 38 | 8239 | 8276 | 84.21 | 0 |
| Chromosome_10 | 4.32 | 15343792 | 15343759 | 34 | 8269 | 8302 | 85.29 | 0 |

| | | | | | | | | |
|---|---|---|---|---|---|---|---|---|
| Chromosome_6 | 3.76E-159 | 11711359 | 11711761 | 403 | 8387 | 8790 | 91.34 | 0.25 |
| Chromosome_3 | 1.01E-20 | 11116217 | 11116305 | 89 | 8429 | 8517 | 86.52 | 0 |
| Chromosome_11 | 4.32 | 9367798 | 9367771 | 28 | 8523 | 8549 | 92.86 | 3.57 |
| Chromosome_10 | 5.98E-49 | 4300259 | 4300605 | 347 | 8582 | 8939 | 73.57 | 7.9 |
| Chromosome_10 | 4.32 | 15800463 | 15800431 | 33 | 8701 | 8735 | 85.71 | 5.71 |
| Chromosome_8 | 6.84E-04 | 10786324 | 10786369 | 46 | 9132 | 9177 | 84.78 | 0 |
| Chromosome_3 | 4.32 | 2177704 | 2177739 | 36 | 9209 | 9246 | 84.21 | 5.26 |
| Chromosome_12 | 4.32 | 2614352 | 2614322 | 31 | 9220 | 9250 | 87.1 | 0 |
| Chromosome_1 | 4.32 | 6452262 | 6452287 | 26 | 9242 | 9267 | 92.31 | 0 |
| Chromosome_5 | 1.24 | 13465092 | 13465060 | 33 | 9323 | 9356 | 88.24 | 2.94 |
| Chromosome_9 | 1.24 | 12021507 | 12021486 | 22 | 9668 | 9689 | 100 | 0 |
| Chromosome_7 | 5.24E-56 | 12063430 | 12063563 | 134 | 10388 | 10521 | 97.01 | 0 |
| Chromosome_3 | 8.89E-28 | 11116431 | 11116623 | 193 | 11307 | 11500 | 75.26 | 0.52 |
| Chromosome_8 | 0.1 | 10533858 | 10533823 | 36 | 11618 | 11651 | 88.89 | 5.56 |
| Chromosome_8 | 4.32 | 2560444 | 2560467 | 24 | 11758 | 11781 | 95.83 | 0 |
| Chromosome_7 | 1.24 | 13125974 | 13126005 | 32 | 12413 | 12446 | 88.24 | 5.88 |
| Chromosome_7 | 0.1 | 10342897 | 10342928 | 32 | 12685 | 12716 | 90.62 | 0 |
| Chromosome_3 | 2.09E-29 | 14423195 | 14423292 | 98 | 13040 | 13137 | 90.82 | 0 |
| Chromosome_3 | 3.31E-52 | 11115979 | 11116216 | 238 | 13060 | 13304 | 79.18 | 2.86 |
| Chromosome_6 | 4.32 | 4484027 | 4484065 | 39 | 13279 | 13317 | 85 | 5 |
| Chromosome_1 | 4.32 | 5637443 | 5637412 | 32 | 13285 | 13317 | 87.88 | 3.03 |
| Chromosome_7 | 4.32 | 15910866 | 15910835 | 32 | 13285 | 13317 | 87.88 | 3.03 |
| Chromosome_3 | 4.32 | 1513798 | 1513833 | 36 | 13819 | 13854 | 83.33 | 0 |
| Chromosome_5 | 0.35 | 11349044 | 11349071 | 28 | 13828 | 13855 | 92.86 | 0 |

**Table S3**. PCR, long range PCR, and Sanger sequencing primers for each *T. californicus* population.

| Usage | Population | Gene | Direction | Sequence (5' to 3') |
|---|---|---|---|---|
| PCR | AB | *COB* | Forward | ATGGTAGCAGGGTTAGCAGGAT[a] |
| | SC | *COB* | Forward | CTATTAGGTGTTTGTCTGGCAACT |
| | SD | *COB* | Forward | CCTTCTGGGAATTTGTTTGGCG |
| | Universal | *COB* | Reverse | ACATADGGYTCTTCHACCGG |
| Long range PCR | AB | *COX1* | Forward | TGAACCGTCTACCCCCCGT |
| | AB | *COB* | Reverse | CACCCCAATTATTCTAAAAAACAAACT |
| | AB | *COB* | Reverse | GCCGTAAAATGAAGCCTTAGAATT |

[a] The 3' end nucleotide mismatches compared to the other population in a cross are underlined.

**Table S4.** PCR reagent kits and PCR cycling conditions for each population.

| | | Population-specific PCR cycling condition | | |
|---|---|---|---|---|
| **Reagent kit** | **PCR Mastermix** | **SC** | **SD** | **AB** |
| Promega Corporation, Cat # M8295 | Default conditions according to manufacturer's instructions | 1) 95 °C 3 min<br>2) 95 °C 30 sec<br>3) 53.5 °C 45 sec<br>4) 72 °C 60 sec<br>5) Repeat 2)-4) 35X<br>6) 72 °C 5 min<br>7) 4 °C hold | 1) 95 °C 3 min<br>2) 95 °C 30 sec<br>3) 55 °C 45 sec<br>4) 72 °C 45 sec<br>5) Repeat 2)-4) 34X<br>6) 72 °C 5 min<br>7) 4 °C hold | 1) 95 °C 3 min<br>2) 95 °C 30 sec<br>3) 55 °C 40 sec<br>4) 72 °C 30 sec<br>5) Repeat 2)-4) 35X<br>6) 72 °C 5 min<br>7) 4 °C hold |
| New England Biolabs, Cat # M0267 | Default conditions according to manufacturer's instructions | 1) 95 °C 3 min<br>2) 95 °C 30 sec<br>3) 54.5 °C 30 sec<br>4) 68 °C 54 sec<br>5) Repeat 2)-4) 30X<br>6) 68 °C 5 min<br>7) 4 °C hold | 1) 95 °C 3 min<br>2) 95 °C 30 sec<br>3) 54 °C 40 sec<br>4) 68 °C 60 sec<br>5) Repeat 2)-4) 35X<br>6) 68 °C 5 min<br>7) 4 °C hold | 1) 95 °C 3 min<br>2) 95 °C 30 sec<br>3) 55.5 °C 35 sec<br>4) 68 °C 45 sec<br>5) Repeat 2)-4) 35X<br>6) 68 °C 5 min<br>7) 4 °C hold |

**Table S5.** SNP variants between the published AB mtDNA genome (Burton et al. 2007, accession DQ917373; Barreto et al. 2018) and the most common AB haplotype.

| Position | Common AB haplotype[a] | Published AB mtDNA |
|---|---|---|
| 1,823.1 | :[b] | A |
| 2,006.1 | : | A |
| 3,976 | G | C |
| 4,706 | T | C |
| 4,707 | G | T |
| 4,708 | A | T |
| 4,744 | T | C |
| 4,761.1 | : | C |
| 4,788 | A | T |
| 4,876 | T | C |
| 5,192 | A | C |
| 5,216 | G | C |
| 5,951 | C | T |
| 6,239 | T | : |
| 6,252.1 | : | T |
| 6,832 | C | T |
| 7,463 | T | C |
| 7,577.1 | : | A |
| 7,580.1 | : | T |
| 7,581.1 | : | A |
| 7,586 | T | C |
| 7,603 | G | A |
| 7,632 | G | A |
| 7,654 | A | C |
| 7,679 | T | C |

| Position | Common AB haplotype | Published AB mtDNA |
|---|---|---|
| 7,700 | T | A |
| 7,708 | G | A |
| 7,710 | G | A |
| 7,720 | T | C |
| 7,747 | T | C |
| 7,778 | T | G |
| 7,788 | T | G |
| 7,847 | T | G |
| 8,900 | T | C |
| 9,517 | A | T |
| 9,520 | A | T |
| 9,524 | A | C |
| 10,102 | T | C |
| 10,112 | T | C |
| 10,204 | C | G |
| 12,099.1 | : | A |
| 12,216.1 | : | A |
| 12,539 | C | A |
| 12,540 | A | C |
| 12,586 | C | A |
| 12,589 | G | C |
| 12,658 | T | C |
| 13,031.1 | : | C |
| 13,400 | C | T |

[a] The common AB haplotype is the consensus sequence of AB Illumina data (NCBI SRA SRX2746703), and was used as the reference for multiple analyses in the present study. All nucleotide positions along the most common AB haplotype are reported here.
[b] The ':' in one haplotype indicates there was an insertion in the other haplotype.

**Figure S1.** Gel image of PCR using potential sources of DNA contamination. Samples used for PCR consisted of relatively clear water, water and algae from populated AB petri dishes, needle tool that contacted an AB individual, and AB individuals all in lysis buffer. Lysis buffer with no DNA, the PCR mastermix with no DNA (-), and empty wells (e) were also included on the gel.

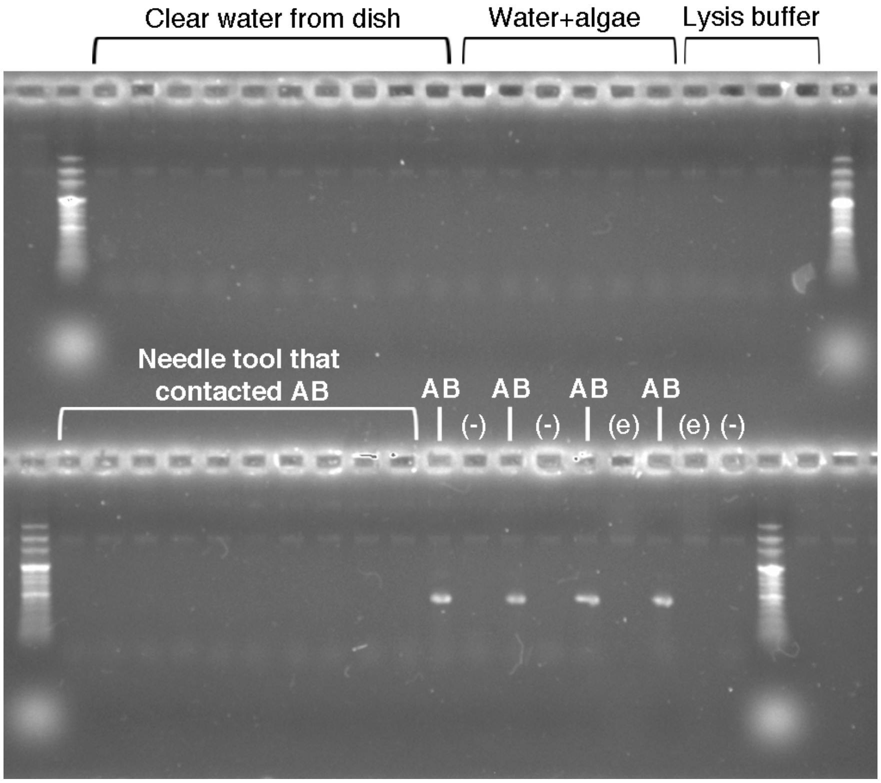

**Figure S2**. Long range PCR products from plasmid-enriched and pooled SC, AB, and SCf x ABm F1 copepods using AB specific primers, in 1% agarose gel stained using ethidium bromide. The first and last lanes contain the entire range of the 1kb DNA ladder (Green BioResearch, catalog # GBR204); the following lanes show DNA from a pool of maternal population SC copepods, paternal population AB copepods, SCf x ABm F1 copepods, and a negative control consisting of the PCR master mix with no DNA. The F1 shows a long paternal AB mtDNA fragment.

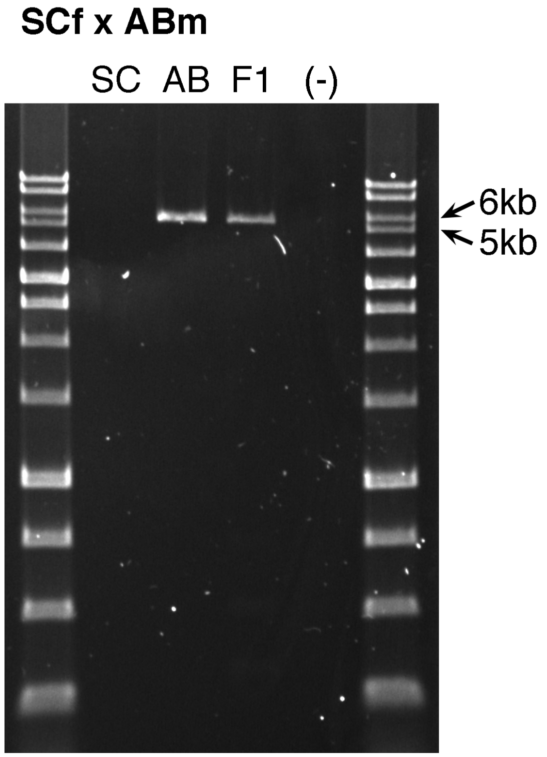


REFERENCES ONLY IN SUPPLEMENTARY MATERIALS


Bankevich A, Nurk S, Antipov D, Gurevich AA, Dvorkin M, Kulikov AS, Lesin VM, Nikolenko SI, Pham S, Prjibelski AD, et al. 2012. SPAdes: a new genome assembly algorithm and its applications to single-cell sequencing. *Journal of computational biology* 19:455-477.

Li H, Durbin R. 2009. Fast and accurate short read alignment with Burrows–Wheeler transform. *bioinformatics* 25:1754-1760.
